# Bacterial small RNAs and host epigenetic effectors of a transgenerational memory of pathogens in *C. elegans*

**DOI:** 10.1101/2021.03.26.437277

**Authors:** Marcela Legüe, Blanca Aguila, Bernardo Pollak, Andrea Calixto

## Abstract

The inheritance of memories is adaptive for survival. Microbes interact with all organisms challenging their immunity and triggering behavioral adaptations. Some of these behaviors induced by bacteria can be inherited although the mechanisms of action are largely unexplored. In this work, we use *C. elegans* and its bacteria to study the transgenerational RNA dynamics of an interspecies crosstalk leading to a heritable behavior. Heritable responses to bacterial pathogens in the nematode include avoidance and pathogen-induced diapause (PIDF), a state of suspended animation to evade the pathogen threat. We identify a small RNA RsmY, involved in quorum sensing from *P. aeruginosa* as required for initiation of PIDF. Histone methyltransferase SET-18/SMYD3 is also needed for PIDF initiation in *C. elegans*. In contrast, SET-25/EHMT2 is necessary for the maintenance of the memory of pathogen exposure in the transgenerational lineage. This work can be a starting point to understanding microbiome-induced inheritance of acquired traits.

## Introduction

Microbes vastly influence life history traits of their host organisms. Some of these traits are adaptive behavioral strategies to microbial pathogenesis that are inherited to the progenies and ensure their survival. How these memories are established and what are the bacterial determinants that trigger them is largely unknown. Bacteria and animals interact persistently, and their relationship has played a crucial role in evolution (Alegado & King, 2014; Provorov, Vorob’ev, & Andronov, 2008). Bacterivore nematodes are among the most ancient and abundant metazoans in the biosphere (Poinar, 2015; Cobb, 1914; van den Hoogen et al., 2019; Samuel, Rowedder, Braendle, Félix, & Ruvkun, 2016). The nematode *Caenorhabditis elegans* feeds on a great variety of bacteria from commensals to human pathogens. The bacteria-*C. elegans holobiont* offers an excellent frame to gain insight into long-term interspecies relationships, and bidirectional phenotypic modulation (Celluzzi & Masotti, 2016; Legüe & Calixto, 2019). *C. elegans* manifests both transient and long-term adaptive behaviors to detrimental bacterial diets. For instance, worms identify pathogens based on their smell and learn to avoid them (Zhang, Lu, & Bargmann, 2005). Attractive or repulsive odors can also generate long-lasting heritable memory (Remy, 2010; Moore, Kaletsky, & Murphy, 2019; Pereira, Gracida, Kagias, & Zhang, 2020). To avoid infection in the absence of food alternatives, part of the progeny of pathogen-fed nematodes undergoes Pathogen-Induced Diapause Formation (PIDF), as a survival strategy for the collective (Palominos et al., 2017; Gabaldón et al., 2020). In this paradigm, pathogen memory is transgenerational, maternally inherited, and requires many effectors of the RNA interference machinery. These properties denote a possible role of *holobiont* small RNAs (sRNAs) in triggering PIDF.

sRNAs can influence multiple generations, either by amplification and maintenance of heritable small RNAs (Houri-Zeevi & Rechavi, 2017) or indirectly, by sRNA-mediated epigenetic modifications (Lev et al., 2017; Kalinava, Ni, Peterman, Chen, & Gu, 2017). Engineered dsRNAs can be transferred between generations in *C. elegans* (Marré, Traver, & Jose, 2016) and moved from distant tissues to the germline (Jose, 2015; Devanapally, Ravikumar, & Jose, 2015; Posner et al., 2019). A question that emerges is whether interspecies sRNAs can modulate life history traits in the hosts that last for generations. In acute host-pathogen interactions bacterial non-coding RNAs can modulate both virulence of bacteria and impact transcription in the host (Westermann et al., 2016; Koeppen et al., 2016; Low et al., 2018; Betin et al., 2019) and host sRNAs can affect bacteria (Liu et al., 2016). A bacterial sRNA influences pathogen avoidance in *C. elegans* (Kaletsky et al., 2020), suggesting this can be one of the mechanisms by which bacteria impact host behavior. We sought to identify the bacterial triggers of PIDF. We identified that RsmY, a bacterial sRNA involved in quorum sensing as a mediator of the defensive behavior in the nematode. Furthermore, we studied histone modifications that could mediate transgenerational PIDF memory. We show that SET-18/ SMYD3, is needed for PIDF to take place. Importantly, SET-25/ EHMT2 is necessary for maintaining the transgenerational memory of pathogen encounter.

In this study, we also dissect the sRNA dynamics in naïve bacteria and in the holobiont throughout their bacteria-host relationship for six generations in a transgenerational paradigm. In this paradigm animals are fed pathogenic bacteria interrupted by two generations feeding on non-pathogens to assess the transgenerational component. We identified the cohorts of sRNAs from pathogenic and non-pathogenic bacteria in naïve and intestinal states. We found that specific bacterial sRNAs change their expression levels between nematode generations. Gene expression also evolves in the worm over each generation, suggesting an intimate reciprocal interaction between bacteria and host. Altogether, our findings provide novel insights into the bacteria-host dynamics across generations, by correlating global sRNA-transcriptomics with an adaptive behavior in *C. elegans*.

## Results

### Bacterial sRNA expression depends on the host previous exposure to same bacteria

The intestinal colonization by pathogenic bacteria triggers a transgenerational defensive behavior in *C. elegans* consisting of microbial exclusion by mouth closure (**Figure 1A, B** and Palominos et al., 2017). In contrast to diapause formation induced by starvation or crowding, pathogen induced diapause is elicited in the second generation after pathogen exposure. Therefore, we explored bacterial sRNAs unique from the second generation that could lead to PIDF. We performed sRNA-seq of bacteria unexposed to host (herein naïve bacteria) and Dual-RNAseq of F1 and F2 nematodes, 24 hours after feeding naïve bacteria from birth (**Figure S1**). Because current sRNA annotations are limited (Diallo & Provost, 2020; Subramanian, Bhasuran, & Natarajan, 2019; Liu et al., 2018), we used an unbiased strategy to find all expressed transcripts. This method finds transcriptional peaks (TPs) and uses their genomic coordinates to define sRNA (Gabaldón et al., 2020). We classified them as *known* sRNAs; *partially novel*; and *novel*, located in intergenic regions (**Figure S2**). Differential expression of bacterial sRNAs was done comparing the transcriptomes of naïve with intestinal bacteria. This experiment allowed us to discriminate gene expression in bacteria in experienced F2 versus inexperienced F1 hosts. Bacterial ncRNAs differentially expressed in F1 and F2 are shown in **Table 1** (Full list of TPs and their genomic contexts is shown in **Table S1)**. Two ncRNA transcripts, the sRNAs RsmY and the tRNA-Gly PA4277.2 were specifically upregulated in the second generation of animals (**Figure 1C**). This suggests that bacteria can sense host previous experience and adjust their sRNA expression in response. RsmZ, an sRNA in the same pathway with RsmY (Valverde, Heeb, Keel, & Haas, 2003), is overexpressed early and stably in the interaction with *C. elegans*. RsmY and RsmZ sRNAs are involved in Quorum Sensing (Kay et al., 2006), a mechanism of bacterial communication (Rutherford & Bassler, 2012). The dynamic expression of RsmY and RsmZ could have a role in interspecies communication leading to behavioral adaptations in the host.

**Figure 1.**
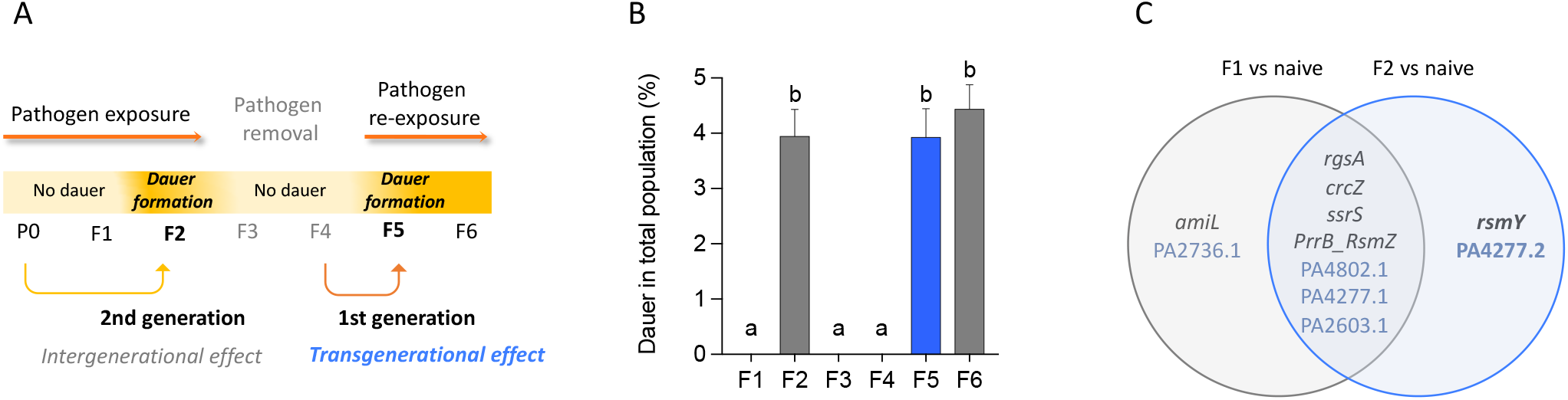
**A**. Schematic representation of the transgenerational paradigm of Pathogen Induced Dauer formation (PIDF), **B**. Quantification of dauer formation for six generation in the transgenerational PIDF paradigm shown in A. a and b denote conditions that are significantly different from each other. condition with the same letter are not significantly different. **C.** Diagram showing the specific expression of *P. aeruginosa* ncRNAs in the intestine of *C. elegans* in the first and the second generation exposed to pathogens. Details of statistical analysis can be found in Dataset 1.

**Table 1.** Differentially expressed genes compared to Naïve.

| Differentially expressed genes compared to Naïve |  |  |  |
| --- | --- | --- | --- |
| F1 |  | F2 |  |
| Gene name | Biotype | Gene name | Biotype |
| *rgsA* | ncRNA | *rgsA* | ncRNA |
| *amiL* |  | *ssrS* |  |
| *crcZ* |  | *crcZ* |  |
| *ssrS* |  | *RsmY* | ncRNA |
| *RsmZ* | sRNA | *RsmZ* | sRNA |
| PA2736.1 | tRNA | PA4802.1 | tRNA |
| PA4802.1 |  | PA4277.2 |  |
| **PA4277.1** |  | **PA4277.1** |  |
| **PA2603.1** |  | **PA2603.1** |  |

### Quorum sensing RsmY sRNA is required for PIDF defensive response

We tested the role of Rsm sRNAs in PIDF by using single and double mutant bacteria of RsmY and RsmZ (Kay et al., 2006) and RsmZ (Heurlier et al., 2004). First, we asked whether rsm mutations retain the ability to infect *C. elegans* by measuring population growth and avoidance behavior. *C. elegans* population size on Δ*rsmY* or Δ*rsmZ* was similar to those grown on wild type *P. aeruginosa* PAO1, while growth in the double mutant Δ*rsmYZ* was larger (**Figure 2A**). Wild type *P. aeruginosa* PAO1 induces lawn avoidance in nematodes 24 after the first exposure (**Figure 2B)**. Avoidance of Δ*rsmY* and Δ*rsmZ* lawns was similar to wild type bacteria. Δ*rsmYZ* however, did not induce avoidance (**Figure 2B**). These results show that single *rsm* mutations do not affect the fitness of pathogens *in vivo* while loss of both genes eliminates virulence. We quantified the growth of each mutant in *C. elegans* intestines by counting colony-forming units (CFU). Δ*rsmY* and Δ*rsmZ* formed over 10^3^ individual bacteria per worm intestine, a large but significantly smaller number than wild type bacteria (Figure 2C). Contrary to single mutants, Δ*rsmYZ* bacteria were many more in number than wild type *in vivo* (**Figure 2C**). This suggests that virulence is not correlated with a greater number of colonies in the intestine, and that the dynamics of organization inside the intestinal lumen may be more relevant for the bacteria-worm communication.

**Figure 2.**
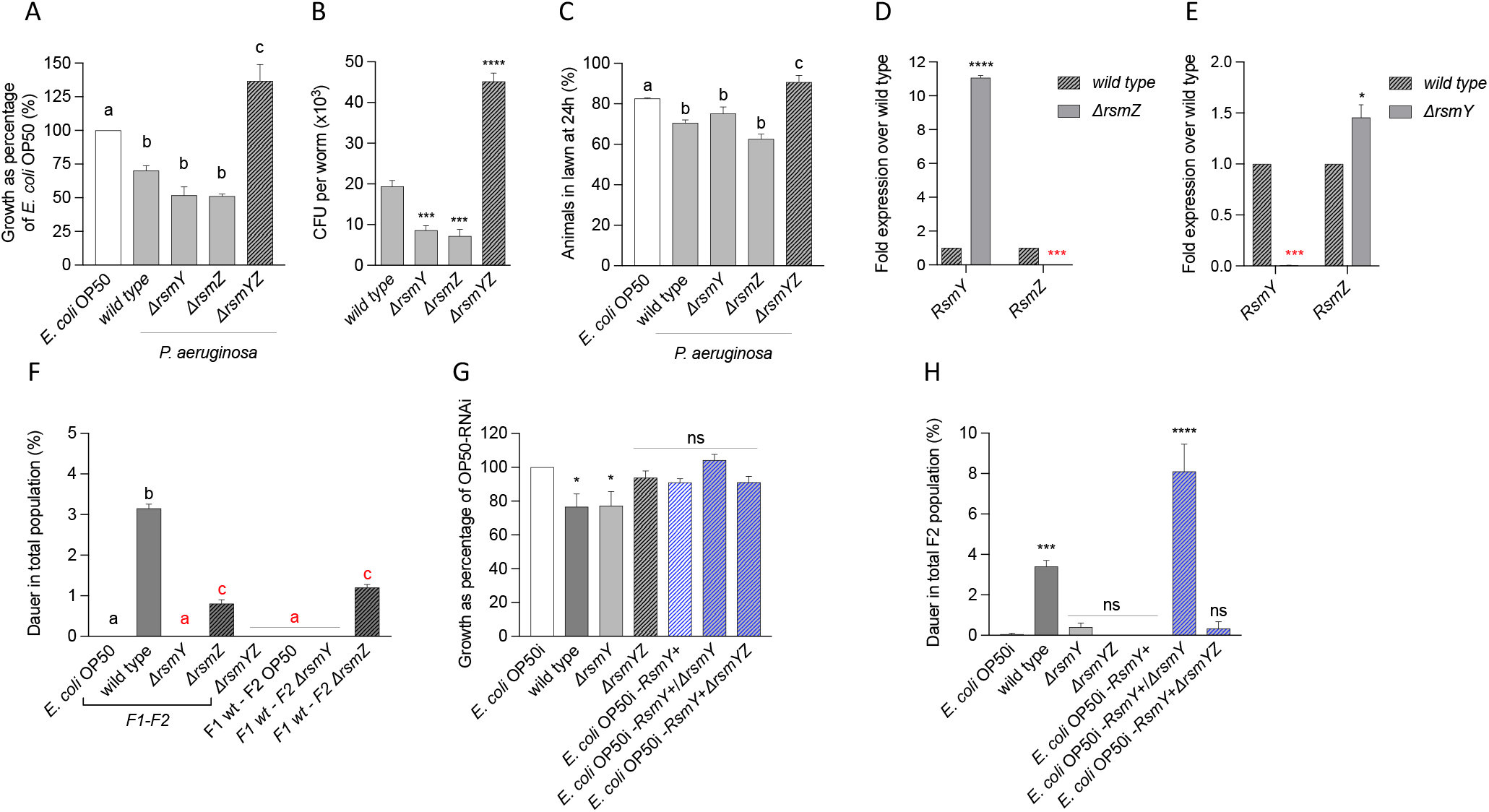
**A**. Quantification of *C. elegans* growth on *rsm* mutants as percentage of growth on *E. coli* OP50. **B**. Quantification of colony-forming units (CFU) in *C. elegans* intestine of wild type, single and double mutants of RsmY and RsmZ. **C.** Quantification of percentage of animals that remain in lawns of wild type, single and double mutants of RsmY and RsmZ. **D**-**E**. Quantification of RsmY and RsmZ RNA expression in Δ*rsmY* and Δ*rsmZ* mutant bacteria. **F**. PIDF quantification for two generations on wild type and *rsm* mutants. **G-H**. Growth (G) and Dauer formation (H) in animals feeding on bacteria expressing RsmY heterologously. In A, C and F, a, b and c denote conditions that are significantly different from each other. Conditions with the same letter are not significantly different. P-value *<0.0332, **<0.0021, ***<0.0002, ****<0.0001.

We grew *C. elegans* in *rsm* lawns for two generations to measure PIDF. Δ*rsmY* failed to induce diapause in the F2, while Δ*rsmZ* induced a smaller percentage than wild type (**Figure 2D**). As expected, avirulent Δ*rsmYZ* did not induce PIDF. To address the requirement of RsmY specifically in the F2, embryos from F1 hermaphrodites grown on wild type *P. aeruginosa* were switched to Δ*rsmY* or Δ*rsmZ* in the second generation. Hypochlorite treatment to obtain embryos between generations eliminates previous bacteria. The change from wild type *P. aeruginosa* to Δ*rsmY* abolished PIDF (**Figure 2D**) while the change to Δ*rsmZ* caused fewer dauers than wild type. These results show that RsmY is necessary for defensive diapause entry in *C. elegans*, with a key role in the F2. RsmZ although not indispensable for PIDF, is required for the wild type response. Δ*rsmY* mutant expressed similar levels of RsmZ than wild type *P. aeruginosa* while Δ*rsmZ* bacteria show over a 10-fold increase in RsmY (**Figure 2E**), consistent with RsmZ being a repressor of the RsmY promoter (Kay et al., 2006). This result also shows that RsmZ cannot compensate the loss of RsmY in triggering PIDF. Taken together, these results show that loss of *rsmY* but not *rsmZ* eliminates defensive diapause in the host.

To test the ability of *rsmY* RNA to induce diapause heterologously we inserted RsmY sequence into a vector used to express double stranded RNA (Fire et al., 1998). An *E. coli* OP50 modified to produce dsRNA (Xiao et al., 2015) was transformed to express RsmY (herein *E. coli-rsmY*+). *E*. *coli-rsmY*+ was induced to produce dsRNA and fed to nematodes (**Figure S3**). *E. coli-rsmY*+ bacteria were unable to trigger PIDF, suggesting that other pathogen’s components are also required for the elicitation of behavior (**Figure 2F**). We fed animals with 80% *E. coli-rsmY*+ mixed with 20% single mutant Δ*rsmY*, or *E*. Δ*rsmYZ*. Δ*rsmY* but not Δ*rsmYZ* mixed with *E. coli-rsmY*+ triggered the production of large numbers of dauers in the second generation while the mutant pathogens alone did not (**Figure 2F**). This result shows that RsmY expression in *E. coli* is sufficient for dauer induction in the presence of a pathogenic background with intact RsmZ. This suggests that RsmZ is needed for pathogen recognition highlighting a different but complementary role for each sRNA in triggering PIDF.

RsmY and RsmZ can promote PIDF by potentially acting on bacterial or *C. elegans* gene expression. We examined *in silico* the existence of perfectly matching sequences to RsmY and RsmZ in the *C. elegans* genome using the BLAST+ tool (Camacho et al., 2009). Our criteria for *bona fide* interaction was an E-value <1. A number of hits, located in coding and non-coding regions (**Table S2**), did not meet the stringency parameters (see Materials and Methods). Our results suggest that if RsmY/Z interact with sequences in *C. elegans* they may use other mechanisms such as imperfect pairing as for miRNAs (Zhao et al., 2017; Gu et al., 2017) or to RNA binding proteins such as in Pseudomonas (Brencic et al., 2009).

### Global sRNA dynamics in *P. aeruginosa* - *C. elegans* interaction across generations

The first time *C. elegans* feeds on *P. aeruginosa* two generations are needed to elicit PIDF. However, PIDF is immediately established after the re-exposure with the pathogen indicating a memory of the first encounter has been transgenerationally inherited. To get insight into the global transcriptomic changes relevant for PIDF memory we performed Dual-small RNA-seq (Westermann, Gorski, & Vogel, 2012) of nematodes and their intestinal bacteria in a transgenerational setting (**Figure S4**). Animals were fed on *P. aeruginosa* for two generations (F1-F2), the F3 was placed on non-pathogenic *E. coli* OP50 until the F4, and the F5 and F6 generations were re-exposed to *P. aeruginosa*.

Changes in sRNAs between naïve and intestinal bacteria as well as throughout the generations in the holobiont, were assessed using the DeSeq2 tool (Love, Huber, & Anders, 2014). sRNA expression of intestinal bacteria was compared to their naïve states (**Figure 3A and B**), and *C. elegans* expression in *P. aeruginosa* was compared to those continuously feeding on *E. coli* OP50 (**Figure 3C**). The intestinal interaction between bacteria and nematode causes large changes in gene expression in the holobiont throughout the generations (**Figure 3D, Files S1-3**). Upon encounter with the host intestine, downregulation of genes is the most frequent change in *bacteria* (**Figure 3E and F**). In *C. elegans* however, the largest response in numbers of genes is upregulatory in the first encounter or re-encounter with the pathogen (F1 and F5, **Figure 3G**). In the subsequent generations (F2 and F6) global changes are much smaller. This suggests the establishment of a gradual adjustment between the two species over the course of interaction.

**Figure 3.**
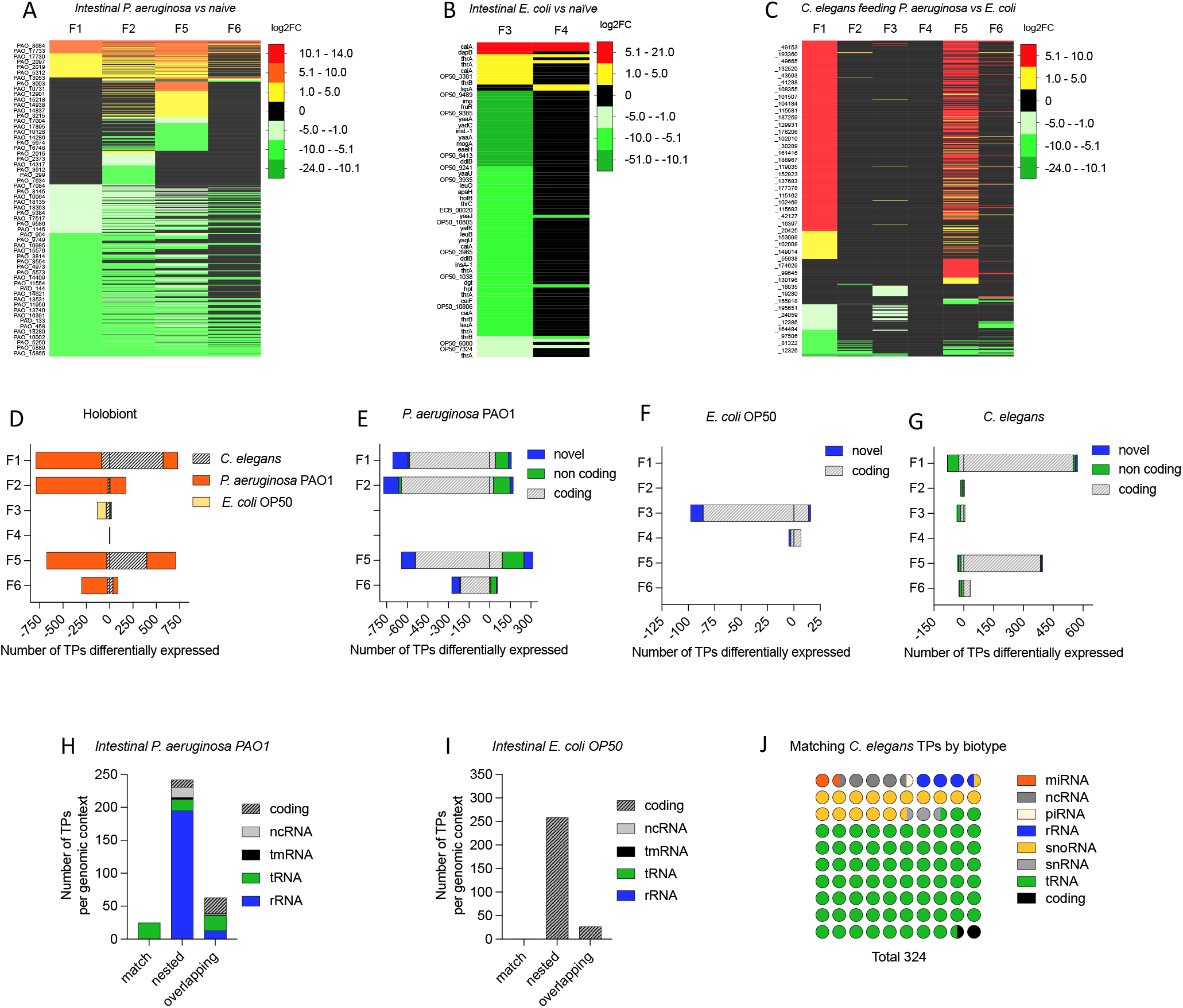
**A-C**. Changes in gene expression in *P. aeruginosa* (**A**), *E. coli* OP50, (**B**) or *C. elegans* (**C**). **D.** Number of TPs differentially expressed in the holobiont (*C. elegans-P. aeruginosa* or *C. elegans-E. coli*) on each generation of interaction. **E-G**. Number of TPs of each naïve bacteria that are matching, nested, or overlapped in coding and non-coding genes in *P. aeruginosa* (**E**), *E. coli* (**F**) and *C. elegans* (**G**). H-I Number of normalized TPs of intestinal *P. aeruginosa* (H) and *E. coli* (I) per genomic context on each RNA biotype. **J**. Number of *C. elegans* TPs expressed in any generation of the transgenerational paradigm per biotype of the known TPs. P-value *<0.0332, **<0.0021, ***<0.0002, ****<0.0001

Host-naïve and intestinal bacteria, and nematodes express a high number of previously unreported TPs. Over 90% of transcripts in all species were either novel or partially novel (**Figure S5**), in coherence with previous reports of extensive pervasive transcription (Lybecker, Bilusic, & Raghavan, 2014; Leitão, Costa, Gabriel, & Enguita, 2020). Most TPs of naïve and intestinal *P. aeruginosa* PAO1 partially matched regions of non-coding RNAs, while all *E. coli* OP50 TPs were nested or overlapped in coding regions (**Figure 3H, I** and **S6A and B**). Known sRNAs in both bacteria and nematodes were mostly tRNAs (**Figures 3H, I and J**). Other biotypes of known transcripts in *C. elegans* include miRNAs, snoRNAs, rRNAs, piRNAs and snRNAs (**Figure 3J**).

We defined as transgenerational sRNAs those TPs that remained differentially expressed after pathogen removal for two generations (F3 and F4). A number of TPs remained overexpressed upon pathogen removal in the F3, but this effect wore off in the F4 (**Figure 3C and G**). This suggests that transgenerational memory of pathogens is likely encoded in other forms of epigenetic memory. Intestinal *E. coli* in the F3 progeny of F2 parents fed with *P. aeruginosa* represses most TPs (**Figure 3F**). Differential expression of *E. coli* in the F4 is almost nonexistent indicating that intestinal *E. coli* becomes transcriptionally similar to the *E. coli* of animals routinely exposed to this bacterium (**Figure 3F**). Altogether these results suggest that the transcriptional repertoires of communicating species are tightly co-dependent.

### Histone modifications are required for PIDF and diapause transgenerational memory

The transgenerational memory of pathogens that induces dauer formation lasts for 5 generations in the absence of pathogens (Palominos et al., 2017). sRNAs and chromatin modifications are effectors of transgenerational inheritance (Rechavi et al., 2014; Houri-Ze’evi & Rechavi, 2016). Because we did not find sRNAs that remained expressed in the F4 generation, we explored whether other transgenerational mediators are needed for PIDF memory after pathogen withdrawal. We tested histone modifications known to participate in transgenerational inheritance such as the histone methyltransferases MET-2, SET-25, SET-32, SET-18 and the chromatin binding protein HPL-2 (Lev et al., 2017; Kalinava et al., 2017). Candidates important for the execution of the dauer decision would fail to enter diapause in the F2 feeding on pathogens. On the other hand, effectors of the transgenerational memory of pathogens would form normal amounts of dauers in the F2 but not in the F5, after re-exposure to pathogenic bacteria. *set-32* and *hpl-2* mutants had impaired growth in non-pathogenic and pathogenic bacteria for two generations, so were unsuitable for testing PIDF (**Figure 4A and B**). *met-2, set-18* and *set-25* mutations supported growth in both bacteria. We quantified the ability of *met-2, set-18* and *set-25* fed *P. aeruginosa* PAO1 to form dauers in the second generation. While *met-2* and *set-25* animals formed normal amounts of dauers, *set-18* animals were unable to do so, suggesting that dimethylation of H3K36 is required for PIDF (**Figure 4C**). To determine the role of *met-2* and *set-25* in transgenerational PIDF, mutants were exposed to pathogens in a transgenerational paradigm (**Figure 1A**). *met-2* animals formed above wild type number of dauers. Interestingly, *set-25* mutants formed normal amounts of dauers in the F2 but failed to form dauers in the F5 (**Figure 4D**), suggesting a role for H3K9 in the transgenerational *memory* of pathogen exposure. Taken together, these results suggest that diverse interspecies players, sRNAs and HMTs contribute to first establish PIDF and posteriorly elicit the memory of the past bacterial encounter in subsequent generations.

**Figure 4.**
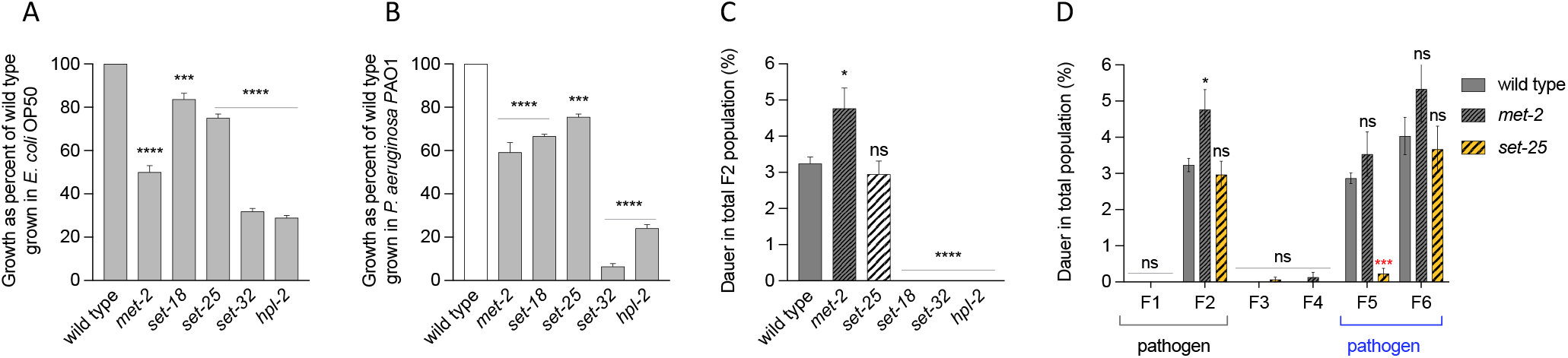
**A-B.** Growth of HMT mutants on *E. coli* (**A**) and *P. aeruginosa* (**B**) as percent of the wild type animals. **C**. Dauer formation of HMT mutants in the second generation. **D**. Transgenerational dauer formation of HMT mutants. P-value *<0.0332, **<0.0021, ***<0.0002, ****<0.0001

## Discussion

Understanding microbiome-induced inheritance is of broad relevance for uncovering the transgenerational origins of health and disease. Here we demonstrate that sRNAs of intestinal pathogens and host histone modifications are required for a transgenerational adaptive behavior in *C. elegans*. Specifically, bacterial *RsmZ* and *RsmY* - both quorum sensing activators - are upregulated sequentially during the colonization of the nematode intestine. *RsmZ* is expressed early and steadily during the interaction while *RsmY* is overexpressed in the second generation of nematodes undergoing pathogenesis. Importantly, RsmY is needed for diapause formation under pathogenesis and its role is key in the second generation. RsmZ however, is not essential but required for the wild type behavioral response. Furthermore, RsmY expressed heterologously promotes PIDF only if RsmZ is present. At the chromatin level the histone methyltransferase SET-18 is necessary for the initiation of the behavior. The trithorax/polycomb protein SET-25 is necessary for the elicitation of the memory of pathogenic encounters. In sum, this work unveils interspecies sRNA dynamics through generations of interaction and histone modifications relevant for the establishment of an adaptive behavioral strategy between interacting species.

### Small RNA dynamics in the holobiont across generations

sRNA expression fluctuated in the holobiont throughout the generations of interactions between nematodes and bacteria. This suggests that a bidirectional regulation of RNA expression is established and depends on the nature of bacteria and the time of residence in the nematode intestine. For example, specific bacterial sRNAs were overexpressed in the second generation of animals suggesting their expression arises from the active co-regulation with the host. We focused on these specific RNAs to find bacterial molecules with a role in triggering the behavioral response that only occurs in the second generation of animals.

In our study, we found a novel role for the bacterial sRNA RsmY in interspecies communication. RsmY is a conserved non-coding RNA in *P. aeruginosa* with a role in communication between bacteria, as quorum sensing (QS) and biofilm regulator (Kay et al., 2006). In *P. aeruginosa*, RsmY and RsmZ are activated by the GacS/GacA system. Both bind and antagonize the RNA binding protein *RsmA*, which negatively controls the expression of quorum sensing and several extracellular products (Pessi et al., 2001; Kay et al., 2006). Our results show that RsmY and RsmZ have different roles in PIDF. We propose that these two ncRNAs act sequentially at successive colonization times. RsmZ is induced early in the interaction with the host, and RsmY expressed in the second generation as a result of sustained interaction. Other Pseudomonas strains capable of inducing behavioral aversion in *C. elegans* also express RsmZ under stress in naive conditions (Kaletsky et al., 2020). We reason these two RNAs could act restricted to the intestine or be internalized to promote an effect in *C. elegans* tissues far from the intestinal lumen. In the first scenario, these sRNAs would impact the lifestyle of intestinal bacteria by affecting the biofilm formation dynamic and therefore impacting intestinal physiology [e.i intestinal distention (Aballay & Hong, 2020)]. In the second scenario, bacterial sRNAs could be internalized through specific dsRNA transporters in the intestine or through outer membrane vesicles (OMVs). By either mechanism sRNAs could reach other organs where direct molecular interactions between bacteria and worm transcripts would occur, using the RNA interference machinery of the host. In this scenario, transcriptional changes could indirectly reach the germline of exposed parents and affect the progeny even when not in direct contact with bacteria (Rechavi & Lev, 2017; Nono et al., 2020; Aballay & Hong, 2020).

These findings broaden our understanding of how small RNAs of colonizing bacteria such as in microbiota regulate host behavior. While host bacterial-RNA targets have been found to promote infection (Koeppen et al., 2016; Zhao et al., 2017), there is scarce evidence about how exogenous sRNA from the microbiota affects physiology and its inheritance in the descendants. This work reveals that bacterial sRNAs are essential for host response to the pathogen. Additionally, bacterial gene expression responds dynamically to changes in gene expression in the host in the different generations. This suggests that bacteria are able to react to the host memory of bacteria exposure, according to a transactional paradigm, in which the *post-interaction* transcriptional state alters the gene expression of upcoming interactions.

### Histone modifications are necessary for initiation of behavior and transgenerational memory

The evidence of microbiome induced phenotypic changes in hosts is increasing, but there is scarce evidence of the long-term epigenetic changes triggered by bacteria. Previously we showed that the interaction with pathogens induces a memory that is inherited transgenerationally (Palominos et al., 2017). This requires that epigenetic information is transmitted by the germline in the absence of direct environmental exposure (Skinner, 2011; Jablonka & Raz, 2009). In our paradigm, the pathogen exposure was maintained for two generations, and then withdrawn for two more generations. The first generation after withdrawal (F3) grew in the uterus of an infected mother, so we could see transcriptomic changes that reflect embryonic exposure. In the next generation (F4) there is no possibility of direct exposure. We did not find either *C. elegans* or bacterial sRNAs that persist in F4, suggesting other transgenerational effectors may play a role in this transmission. In our paradigm, the transgenerational memory is observed after re-exposure, where animals form dauers immediately after the new exposure to pathogens in contrast with the two generations needed the first time animals are fed pathogens. We found that SET-25 is necessary for either storage or retrieval of the inherited memory, even though it is dispensable for intergenerational dauer formation. SET-25 was previously reported to be necessary for the initiation but dispensable for transgenerational maintenance of RNAi silencing (Woodhouse & Ashe, 2019). On the other hand, H3K9me3 is not required for nuclear RNAi-induced transcriptional silencing (Kalinava et al., 2017; Mao et al., 2015). This suggests that the participation of H3K9me3 in transgenerational inheritance may vary depending on the genes and the phenomena studied.

## Materials and Methods

### *C. elegans* maintenance and growth

Wild type (N2) animals and mutant strains MT17463 [*set-25(n5021)*], VC967 [*set-32(ok1457)*], VC767 [*set-18(gk334)*], MT13293 [*met-2(n4256)*], and RB1090 [*hpl-2(ok1061)*] were grown at 20°C as previously described (Brenner, 1974). All nematode strains were maintained on *E. coli* OP50 strain prior to feeding with other bacteria.

### Bacterial growth

The following bacterial strains were used as *C. elegans* food: *E. coli* OP50, RNAi-competent *E. coli* rnc14::DTn10 and lacZgA::T7pol camFRT, *P. aeruginosa* PAO1, PAO6354 (Δ*rsmZ*), PAO6420 (Δ*rsmY*) and PAO6421 (Δ*rsmY rsmZ*). Bacteria were grown overnight on Luria-Bertani (LB) plates at 37°C from glycerol stocks. The next morning a scoop of the bacterial lawn was inoculated in LB broth with antibiotic (streptomycin for *E. coli* OP50 and grown for 6 hours on agitation at 450 g at 37°C (OD_600_ 1.5-2.0). A volume of 3 mL of this bacterial culture was seeded onto 90 mm NGM plates and allowed to dry overnight before nematodes are placed on them.

### Quantification of dauer formation

#### In the second generation of pathogen exposure

Quantification of dauers was done as previously reported (Palominos et al., 2017). Entire worm populations on each plate were collected in 1 mL of M9. The initial stock was diluted 1:10 in M9. The total population including dauer larvae was treated with 1% SDS for 20 minutes (Cassada & Russell, 1975). A volume of 10μL of each dilution was used to count the total population and dauer larvae respectively under a Nikon SMZ745 stereomicroscope. Each technical replica was scored 3 times and the number of dauers was plotted as percent of the total population of animals.

#### Transgenerational dauer formation

Quantification of dauers was done as previously reported (Palominos et al., 2017; Chávez & Calixto, 2019) with minor variations. Control animals were maintained on *E. coli* OP50 for six generations, while test animals were exposed to *P. aeruginosa* PAO1 for two generations, passed as F3 embryos to *E. coli* OP50 until the F4, and reintroduced to *P. aeruginosa* PAO1 in the F5 and F6 generations. In detail, we exposed 15 L4 animals (P0) to *P. aeruginosa* PAO1 or *E. coli* OP50 on 90 mm plates. 24 hours later, gravid adults were isolated and treated with sodium hypochlorite (Stiernagle, 2006). As a result, near 6000 clean embryos were deposited in new Petri dishes. This process was made every three days in animals fed on *E. coli* OP50 and every four days on *P. aeruginosa* PAO1. F2 adults were treated with sodium hypochlorite, and the embryos were passed to new plates of *E. coli* OP50. The same treatment was done with the F3 and the F4 generations. F5 embryos from bleached F4 adults were passed to pathogens, and their F6 progeny as well, using the same treatments. Embryos on *P. aeruginosa* PAO1 were allowed to grow for 48 −72 hours before treating the entire plate with 1% SDS for the quantification of dauers. All experiments were done in three biological replicates of three technical replicates each. Importantly, six plates were used for each condition; three were used for dauer counts and three for hypochlorite treatment of adults.

### RNA extraction from bacteria

Bacterial inoculums of *E. coli* OP50, *P. aeruginosa* PAO1, PAO6354 (Δ*rsmZ*), PAO6420 (Δ*rsmY*) and PAO6421 (Δ*rsmY rsmZ*) were taken from glycerol stocks kept at −80°C and grown overnight on LB agar plates at 37°C (with streptomycin 100 μg/mL or without antibiotic, depending on the bacterial strain). Next morning, a scoop of the bacterial lawn was inoculated in 15 mL of liquid LB with the same antibiotics, at 37°C in agitation for 4 hours or until they reached an OD_600_ between 1.5-2.0. A volume of 1.5 mL of the sample was pelletized and incubated with 200 μL of preheated TRIzol^™^ Max^™^ Bacterial Enhancement Reagent (Thermo-Fisher) at 95°C for 4 minutes. After that 1 mL of TRI-reagent (MRC) was added to the pellet and one steel bead the 0.5 mm was added to the tube. Maximal vortex was applied for 5 minutes; the sample was centrifuged and changed to a new tube. Phase separation was performed adding 200 μL of chloroform and vortexing. After centrifugation at 12,000 g for 15 min at 4°C, 400 μL of aqueous RNA phase was precipitated at room temperature overnight, followed by centrifugation for 15 minutes at 4°C. The supernatant was discarded, and the pellet was washed with EtOH 75%, air-dried for 10 minutes and resuspended in 20 μL of DNAse-free water. DNase I treatment was performed adding 2 μL of buffer, 2U of DNAse I, and incubated at 37°C for 10 minutes. Inactivation of DNAse was done with addition of 2 μL of 50 mM EDTA. Each procedure was performed in triplicates. Control of sizing, quantification and quality was made with Agilent 2100 Bioanalyzer, using RNA 6000 Pico kit.

### RNA extraction of *C. elegans* and its intestinal bacteria

Wild type *C. elegans* were cultured on standard 60 mm diameter Petri dishes with NGM media, seeded with 0.5 mL of *E. coli* OP50 and maintained at 20°C. After three days, a decontaminant treatment with NaOH and sodium hypochlorite was applied (Stiernagle, 2006). As result, only clean embryos were deposited in new dishes. After 48 hours, most individuals were in L4 stage. 20 to 25 L4 worms were transferred to 90 mm NGM plates previously seeded with 3 mL of *E. coli* OP50 or *P. aeruginosa* PAO1. In all cases the bacterial lawn covers the plate. Worms were allowed to grow at 20°C for 48 hours. The resulting population was collected with an M9 buffer and treated with sodium hypochlorite to collect only embryos (F1) that were allowed to grow for 24 hours under the same conditions as their progenitors (P0). Half of the F1 larvae was washed with M9 ten times to avoid external bacteria from the worm body, and frozen at −80°C for posterior RNA extraction. The other half was allowed to grow under control or pathogenic conditions for 2 more days (F1 adults). F2 embryos were obtained and grown exactly as the F1. L1-L2 larvae were harvested with an M9 buffer, washed with M9 ten times to avoid external bacteria from the worm body, pelletized and frozen at −80°C for RNA extraction. Each condition was performed in triplicates. Frozen samples of L1-L2 larvae from F1 and F2 are pre-processed for mechanical lysis by vortexing the sample with 0.5 mm steel beads for five minutes. RNA purification from pelleted worms is performed using phenol-chloroform extraction (TRI reagent^®^, Sigma-Aldrich) with the same protocol described above. Our experiment consisted of 18 samples: 6 RNA samples from worms grown on *E. coli* OP50 or *P. aeruginosa* PAO1. For each bacterium 3 samples were from the F1 and 3 from the F2 generations. Quality tests were performed by Nanodrop, Denaturing gel electrophoresis and finally using DNA/RNA 1000 chip kit in a Bioanalyzer 2100 (Agilent Technologies) and quantified by Quant-iT^™^ RiboGreen^®^ Assay kit (Life Technologies).

### Small RNA Sequencing

*C. elegans* and bacterial small RNAs were sequenced using the MiSeq Illumina platform in Centro de Genómica y Bioinformática (CGB), Universidad Mayor. cDNA library preparation was made with Diagenode CATS Small RNA sequencing kit for Illumina ^®^ according to manufacturer’s protocol. All RNA-seq experiments generated Illumina reads in FASTQ format. Raw data is available in NCBI as Bioproject number PRJNA708299.

### Bioinformatic analysis

All the scripts used in the analysis are available at https://bitbucket.org/mlegue/workspace/projects/TSI Data pre-processing and quality control. Quality visualization was made with FastQC (https://www.bioinformatics.babraham.ac.uk/projects/fastqc/). Trimming was performed with Cutadapt (Martin, 2011), with Diagenode recommended executable CATS_trimming_r1.sh for CATS Library Preparation Kits available at https://www.diagenode.com/en/documents/diagenode-trimming-tools-for-cats-rnaseq

Mapping. For each sample, reads were mapped to *E. coli* OP50 genome assembly ASM435501v1, *P. aeruginosa* PAO1.

ASM676v1 versions or *C. elegans* WS267. Alignment was performed with Bowtie2 version 2.2.6 (Langmead & Salzberg, 2012) with one allowed mismatch and seed set to 17 bp. As a result, a bam file was produced for each sample.

We performed an annotation strategy to detect both known and unannotated transcripts from worm and bacteria, based on transcriptional peaks (TPs) coordinates (Gabaldón et al., 2020). The features obtained for both strands were gathered and sorted to create a gff file for further analysis. We kept features between 17 and 150 nucleotides, with an average coverage of 10 reads by nucleotide.

Comparison with annotated genes. We categorize our TPs as known (matching 85% or more of an annotated feature), novel (in intergenic regions), and partially novel (nested or overlapped to annotated features, with less and more than 15% of unmatched nucleotides respectively). All of them are also categorized as sense or antisense to a known reference feature. To accomplish that, we compared our TPs against the Ensembl annotations of *C. elegans* PRJNA13758, WS267 by using the intersect function of BEDTools (Quinlan & Hall, 2010), adapted from Gabaldon et al. (Gabaldón et al., 2020). We performed the same process for *E. coli* OP50 ASM435501v1 and *P. aeruginosa* PAO1 ASM676v1 (Quinlan & Hall, 2010). We also created a TPs fasta file with the feature sequences by using getfasta BEDtools command.

### Differential expression analysis

For each sample read count was performed with featureCounts (Liao, Smyth, & Shi, 2014) from bioconductor Rsubread package with default parameters (Anders, Pyl, & Huber, 2015). Then, the count matrix was used to perform differential expression analysis in R version 3.3.2 (Anders & Huber, 2010) (Robinson, McCarthy, & Smyth, 2010) between the naïve and intestinal bacteria of each generation using DeSeq2 (Love et al., 2014). For analysis in which differential expression is not involved, we generate normalized expression by applying the TMM method (Trimmed Mean of M, Robinson and (Robinson & Oshlack, 2010).

### RsmY crosskingdom matching sequences

We searched for similar sequences between rsmY and a database generated with the *C. elegans* genome (WS267) using Blast + (Camacho et al., 2009).

### RNA extraction in a transgenerational paradigm

60 L4 worms (P0) were placed on plates seeded with *P. aeruginosa* PAO1. After 24 hours the plates were carefully washed with M9 to eliminate larvae and adult worms. The laid eggs (F1) were allowed to grow for 24 hours in *P. aeruginosa* PAO1. F1 L2s were collected with M9 in an Eppendorf tube and washed 5 times with 1 mL of M9 and centrifugation at 2000 rpm. The L2 F1 worms were subjected to total RNA extraction with TRiZol reagent (Invitrogen).

To obtain F2 worms, 20 L4 (P0) worms were seeded on a plate with *P. aeruginosa* PAO1 and allowed to grow for two generations for 7 days until the first F2 embryos were laid. F1 gravid hermaphrodites were treated with hypochlorite and F2 embryos were placed on a plate with fresh PAO1 and allowed to hatch and grow for 24 hours. L2 animals were collected for RNA extraction. A portion of the F2 embryos was placed in a *P. aeruginosa* PAO1 plate until worms reached adulthood. Animals were treated with hypochlorite and their F3 was placed on a plate with *E. coli* OP50. After 24 hours, worms were collected for RNA extraction. A part of F3 embryos were maintained in *E. coli* OP50 until adulthood. F4 embryos were obtained by hypochlorite treatment and allowed to grow for 24 hours when L2 were collected for RNA extraction. A part of F4 embryos were maintained in *E. coli* OP50 until adulthood. Gravid hermaphrodites were then treated with hypochlorite solution and their eggs placed on a fresh plate with *P. aeruginosa* PAO1 for 24 hours. L2 worms were washed, and their total RNA extracted. A part of F5 embryos was placed in a plate with *P. aeruginosa* PAO1 until obtaining embryos of the sixth generation. F5 hermaphrodites were treated with hypochlorite and the F6 embryos were placed on fresh *P. aeruginosa* PAO1 plate for 24 hours. 24 hours later, L2 animals were collected for extraction of total RNA.

### Pathogen avoidance

60 mm NGM plates were seeded with 400 μL of bacteria inoculum incubated at 37°C in agitation for 4 hours. Next day, 30 N2 worms were transferred by picking and placed in the center of the lawn. The number of worms inside and outside the lawn was quantified at 10 and 30 minutes, 1 hour, 2 hours and 24 hours.

### Quantification of intestinal Colony Forming Units

Thirty L4 animals were picked into an Eppendorf tube containing M9 buffer with 25 mM levamisole hydrochloride (Sigma-Aldrich) to cause paralysis and stop pharyngeal pumping (Kawli & Tan, 2008). The animals were then washed three times with M9 containing 1 mg/mL gentamicin (Sigma-Aldrich) and 1 mg/mL ampicillin (Sigma-Aldrich). After the third wash, the animals were incubated once more with the antibiotic mixture for 1 hour. To eliminate the antibiotic, the animals were washed three more times with M9 containing 25 mM levamisole. Each worm pellet was lysed with an individual pestle, and the resulting lysate was serially diluted 1:10 seven times in M9. Amounts of 200 of each dilution were individually plated on LB with streptomycin (to select *E. coli* OP50), and without antibiotics for *P. aeruginosa* PAO1. The plated dilutions were incubated overnight at 37°C. The amount of CFUs was calculated using the following formula: CFU per worm /[(colonies per plate/dilution factor) / plated volume]/number of worms.

### Reverse transcription of bacterial RNA

Two μg of total RNA were treated with DNase I (Invitrogen) according to manufacturer’s instructions. ImProm-II Reverse Transcriptase (Promega) was used to synthesize cDNA from total RNA. Briefly, 1 μg of total RNA with 0.5 μg of random primers (Promega) and distilled water up to 10 μl of final volume were incubated at 70°C for 5 minutes and then kept on ice for 5 minutes. The reverse transcription mix was prepared with 4 μl ImProm-II 5X Reaction Buffer, 2.5 mM MgCl2, 0.75 mM dNTP mix and 1 μl of SuperScript IV Reverse Transcriptase (Thermo Fisher Scientific) for each sample and mixed with 10 μl of the previous reaction. Synthesis was performed in an Aeris Thermal Cycler using the following parameters: annealing at 25°C for 5 min, extension at 42°C for 60 minutes, inactivation of the reverse transcriptase at 70 ° C for 15 minutes.

### Confirmation of mutations in single and double mutants of *P. aeruginosa* PAO1 by PCR

To confirm RsmY and RsmZ expression in the mutant strains PAO6354 (Δ*rsmZ*), PAO6420 (Δ*rsmY*) and PAO6421 (Δ*rsmY rsmZ*), we used the following primers: For amplification of rsmY 5’ AGGACATTGCGCAGGAAG 3’ and 5’ GGGGTTTTGCAGACCTCTATC 3’; for rsmZ 5’ CGTACAGGGAACACGCAAC 3’, and 5’ TATTACCCCGCCCACTCTTC 3’. These primers were designed based on conserved regions of genes in *Pseudomonas aeruginosa* PAO1 (GCF_000006765.1, *Pseudomonas* Genome Database (Winsor et al., 2016). PCR reaction mixtures consisted of 10 μL of KAPA HiFi HotStart ReadyMix 2X PCR Kit (Kapa Biosystems), 0.3 μM forward primer, 0.3 μM reverse primer, 100 ng of bacterial cDNA and water up to 20 μL of final volume. Amplification was performed in an Aeris thermal cycler using the following parameters: initial denaturation at 95°C for 3 min, 98°C for 20 seconds for each subsequent cycle; annealing 58°C for 15 seconds and elongation at 72°C for 15 seconds for 30 cycles; and a final elongation at 72°C for 1 minute.

Amplicons from the PCR amplifications were purified using a QIAquick PCR Purification Kit (Qiagen, Cat No.28104). Sequencing of the PCR product was performed by Capillary Electrophoresis Sequencing (CES) for difficult samples (Macrogen Korea) using the same primers described before.

### RsmY and RsmZ expression quantification in bacteria by Real Time PCR

RsmY and RsmZ expression quantification was done by RT-PCR using Mir-X miRNA qRT-PCR TB Green Kit (Takara Bio USA, Inc). For each reaction, 50 ng of bacterial cDNA, 12.5 μl of TB Green Advantage qPCR 2X, 0.5 μl ROX (50X) LSR, 0.2 μM primers described before (For amplification of rsmY 5’ AGGACATTGCGCAGGAAG 3’ and 5’ GGGGTTTTGCAGACCTCTATC 3’; for rsmZ 5’ CGTACAGGGAACACGCAAC 3’, and 5’ TATTACCCCGCCCACTCTTC 3’.For amplification of rsmY 5’ AGGACATTGCGCAGGAAG 3’ and 5’ GGGGTTTTGCAGACCTCTATC 3’; for rsmZ 5’ CGTACAGGGAACACGCAAC 3’, and 5’ TATTACCCCGCCCACTCTTC 3’)., and water up to 25 μl final volume were used. The StepOnePlus™ Real-Time PCR System was used with the following conditions: Initial denaturation at 95°C for 3 minutes, 30 cycles of 98°C for 20 seconds, 58°C for 15 seconds and 72°C for 15 seconds, followed by a final extension at 72°C for 1 minute.

#### Bacterial DNA fragment amplification

DNA extraction of *P. aeruginosa* PAO1 was performed using UltraClean ^®^ Microbial DNA Isolation Kit (Mo Bio Laboratories Inc.) according to the manufacturer’s instructions. Amplifications of *rsmY* and *rsmZ* genes were performed by PCR (same PCR conditions as previously mentioned in confirmation of mutant strains section) using genomic DNA as template. PCR products were run on 2% agarose gel electrophoresis. Thereafter, fragments were purified using the QIAquick PCR Purification Kit (Qiagen, Cat No.28104) and quantified in Infinite 200 PRO NanoQuant. Finally, DNA fragments were stored at a concentration of 16.66 fmol/μL at −20°C until used.

Construction of L4440-rsmY and L4440-rsmZ

A 500 ng mass of L4440 vector (Fire et al., 1998) was digested using EcoRV (NEB Cat. No. R0195S) and dephosphorylated using Antarctic Phosphatase (NEB Cat. No. M0289S) according to manufacturer’s instructions to generate a linearized blunt-ended fragment for cloning. RsmY and RsmZ were amplified by PCR using oligonucleotides containing flanking SapI restriction enzyme sites. For amplification of rsmY,5’AAGCTCTTCAGGTGTCAGGACATTGCGCAGG3’ and 5’TTGCTCTTCTGTAAAAACCCCGCCTTTTGG3’ were used, and for amplification of rsmZ 5’ AAGCTCTTCAGGTCGTACAGGGAACACGCAAC3’ and 5’TTGCTCTTCTGTAAAAAAGGGGCGGGGTATTAC3’ were used. PCR amplicons were gel purified using the Wizard^®^ SV Gel and PCR Clean-Up System (Promega Cat. No. A9281). Next, a mass of 200 ng of the amplicons was digested using SapI (NEB), followed by a single elongation step PCR by adding an equal volume of previously prepared Phusion polymerase PCR reaction mix at 2X final concentration (Phusion™ High-Fidelity DNA Polymerase, ThermoFisher Cat. No F-530S) without oligonucleotides. This provided 5’ phosphate-containing blunt fragments for cloning. 5’ phosphate-containing rsmY and rsmZ amplicons were then ligated to the linearized dephosphorylated L4440 vector using a 2-hour room temperature incubation with T4 DNA ligase (NEB Cat. No. M0202S) according to the manufacturer’s instructions. The samples were then transformed into TOP10 chemically competent cells and plated onto LB agar plates containing appropriate antibiotics. Resulting colonies were screened by colony PCR, purified by column-purification (Qiagen) and verified by Sanger sequencing (Macrogen Inc., Korea).

### Bacterial transformation

A volume of 100 μL of chemically competent TOP10 cells (Thermo Fisher Scientific), RNAi-competent *E. coli* rnc14::DTn10 and lacZgA::T7pol camFRT and PAO6420 (*ΔrsmY*) were transformed with 2 μL of construct obtained and kept on ice. After 30 minutes of incubation, heat-shock was performed for 45 seconds at 42°C and immediately kept on ice for 5 minutes. Afterwards, 900 μl of LB medium are added and incubated at 37 °C for 2 hours at 200 rpm. Then, 50 μL of each reaction was plated on LB agar plates with 100 μg/mL ampicillin, 40 μg/mL 5-bromo-4-chloro-3-indolyl-beta-D-galactopyranoside (X-Gal) (Thermo Scientific) and 1 mM isopropyl beta-D-thiogalactopyranoside (IPTG, US Biological). Plates were incubated at 37°C for 24 hours. To bacterial colony screening, two white positive colonies were selected for each plate and grown in 5 ml of LB medium with 100 μg /mL of ampicillin and incubated at 37°C for 24 hours.

TOP10 cells positive colonies were extracted using QIAprep Spin Miniprep kit and sequenced at Macrogen Korea. Additionally, constructs were also digested with BsaI enzyme (New England Biolabs), using 1 μL Cutsmart buffer 10x, 0.2 μL BsaI (10U / μL), 100 ng of plasmid and water up to 10 μL final volume. Reactions were incubated at 37 ° C for 1 hour. Subsequently, the reaction was run on 1.5% agarose gel electrophoresis to verify the presence of fragments.

RNAi-competent *E. coli* rnc14::DTn10 and lacZgA::T7pol camFRT and PAO6420 (Δ*rsmY*) positive colonies, total RNA were extracted with TRiZol reagent (Invitrogen), reverse transcription of bacterial RNA and finally confirmation of *rsmY* amplification by PCR previously describe above.

#### Criteria for data exclusion

We excluded experimental replicas when there was contamination with unwanted bacteria or fungi on the nematode plates, or when bacteria have been almost or completely consumed.

##### Experimental replicas and statistic evaluation

All experiments were done at least 3 times (three biological replicas, started in different days and from different starting plates). Each biological replica contained a triplicate (three technical replicas). Statistical evaluation was done by a one or two-way ANOVA with post-hoc analyses. Results of all tests are detailed in **Source Data File 1**.

## Supporting information

SourceDataFile1

## Supplementary Figures

**Figure S1.**
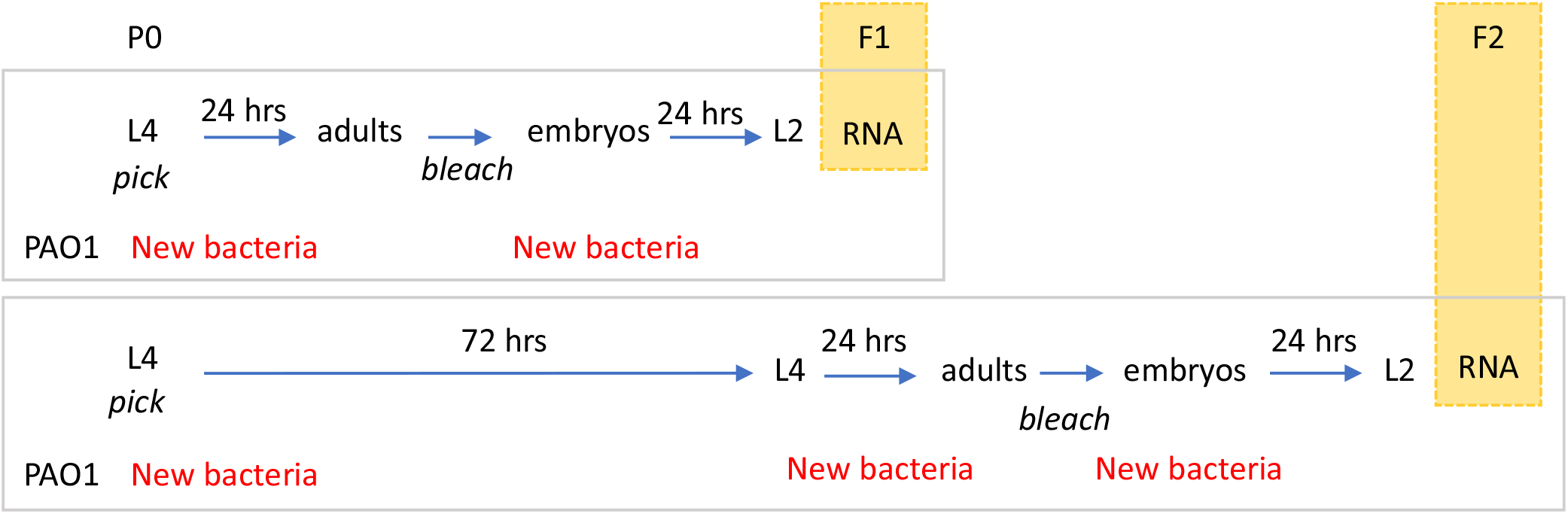
Dual-small RNA-seq of bacteria and worm for two *C. elegans* generations.

**Figure S2.**
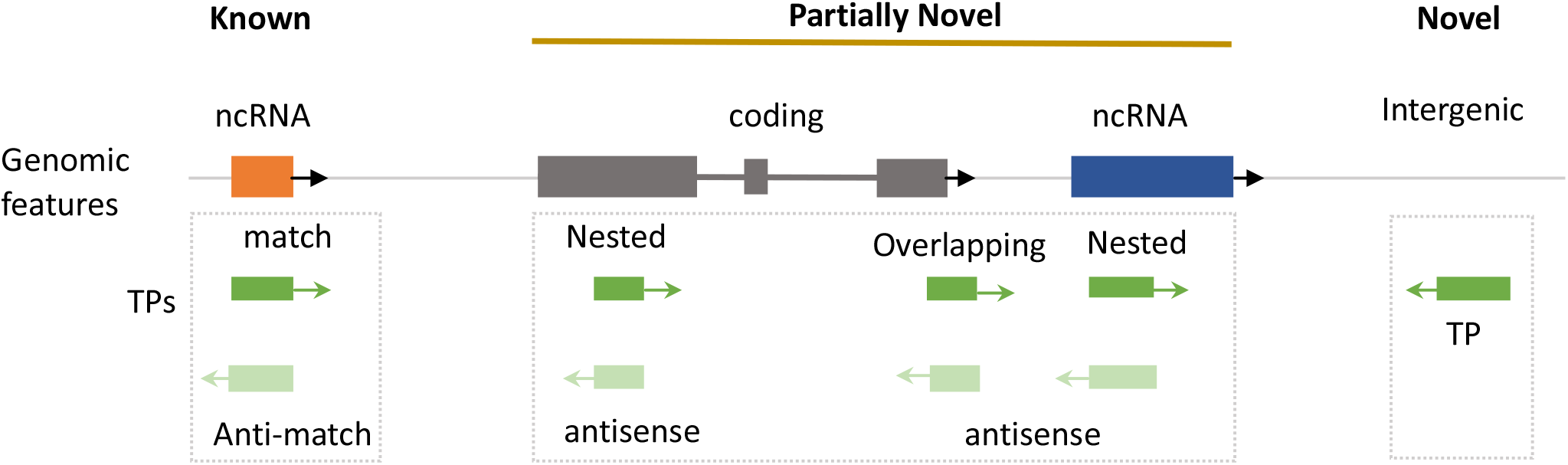
Representation of genomic context of transcriptional peaks (TPs). We classified them as *known* (matching an annotated feature); *partially novel* (nested or overlapping to an annotation); and *novel*, located in intergenic regions.

**Figure S3.**
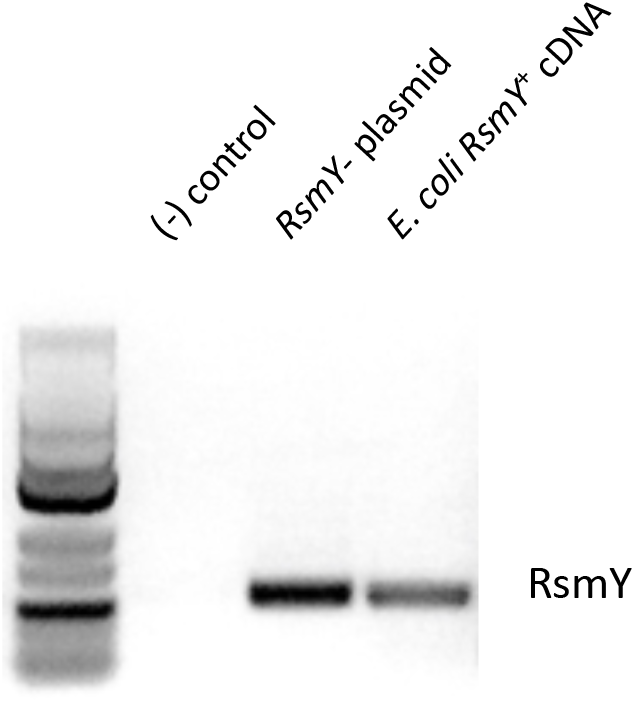
Expression of RsmY sRNA in *E. coli* RsmY+ strain. PCR amplification using RsmY primers. From left to right: Low range RNA ladder, negative control, RsmY+ plasmid, cDNA from *E. coli-* RsmY+.

**Figure S4.**
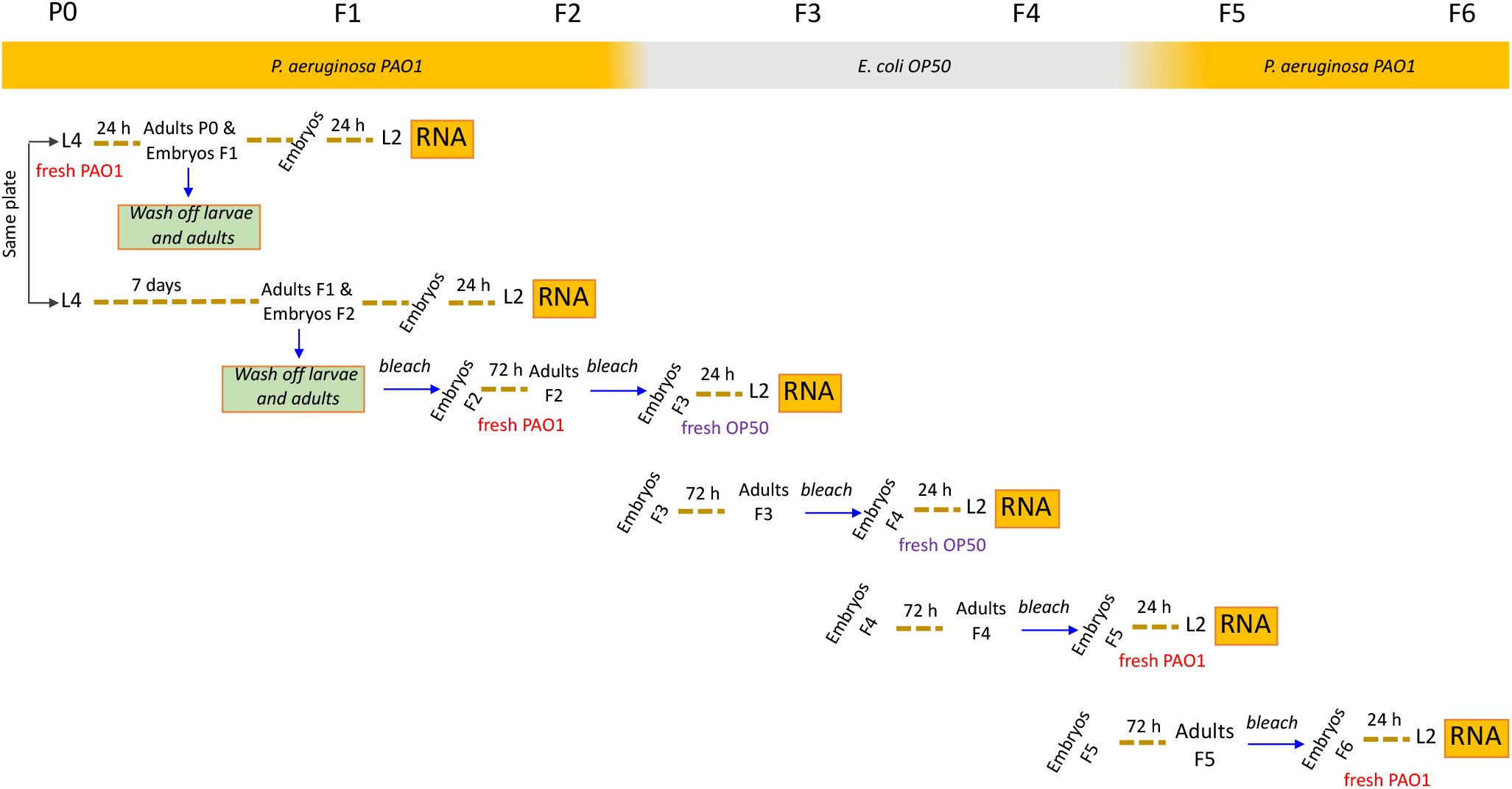
Dual-small RNA-seq of bacteria and worm for six *C. elegans* generations

**Figure S5.**
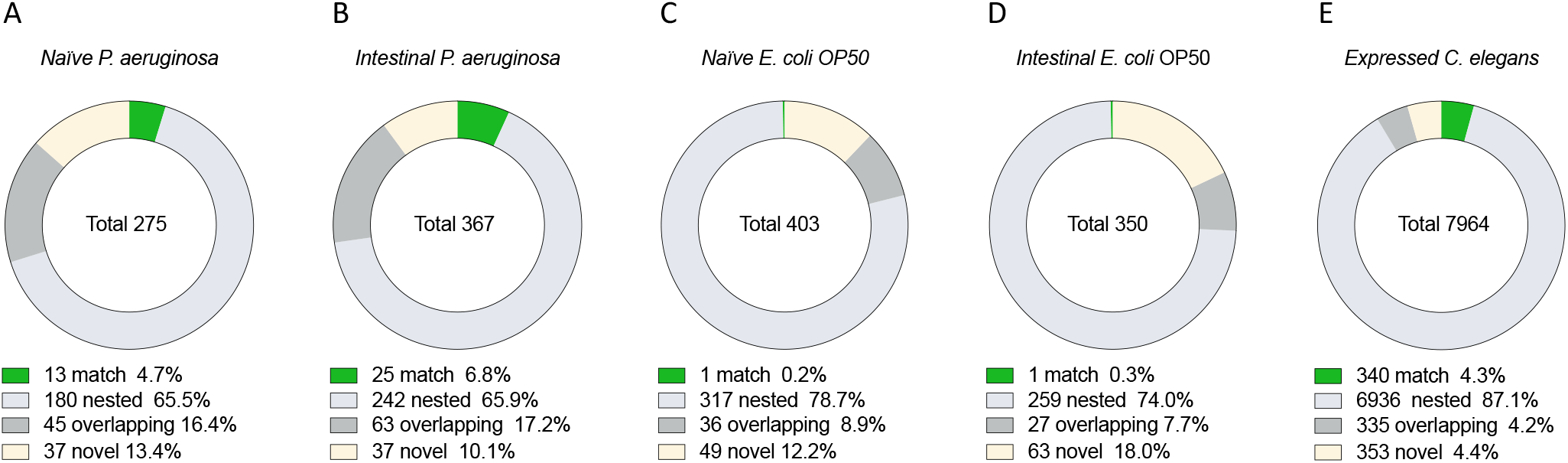
**A-E.** Number of normalized TPs of naïve (A,C) and intestinal (B,D) *P. aeruginosa* (A,B) and *E. coli* (C, D) in each genomic context. E. Number of *C. elegans* TPs expressed in any generation of the transgenerational paradigm per genomic context.

**Figure S6.**
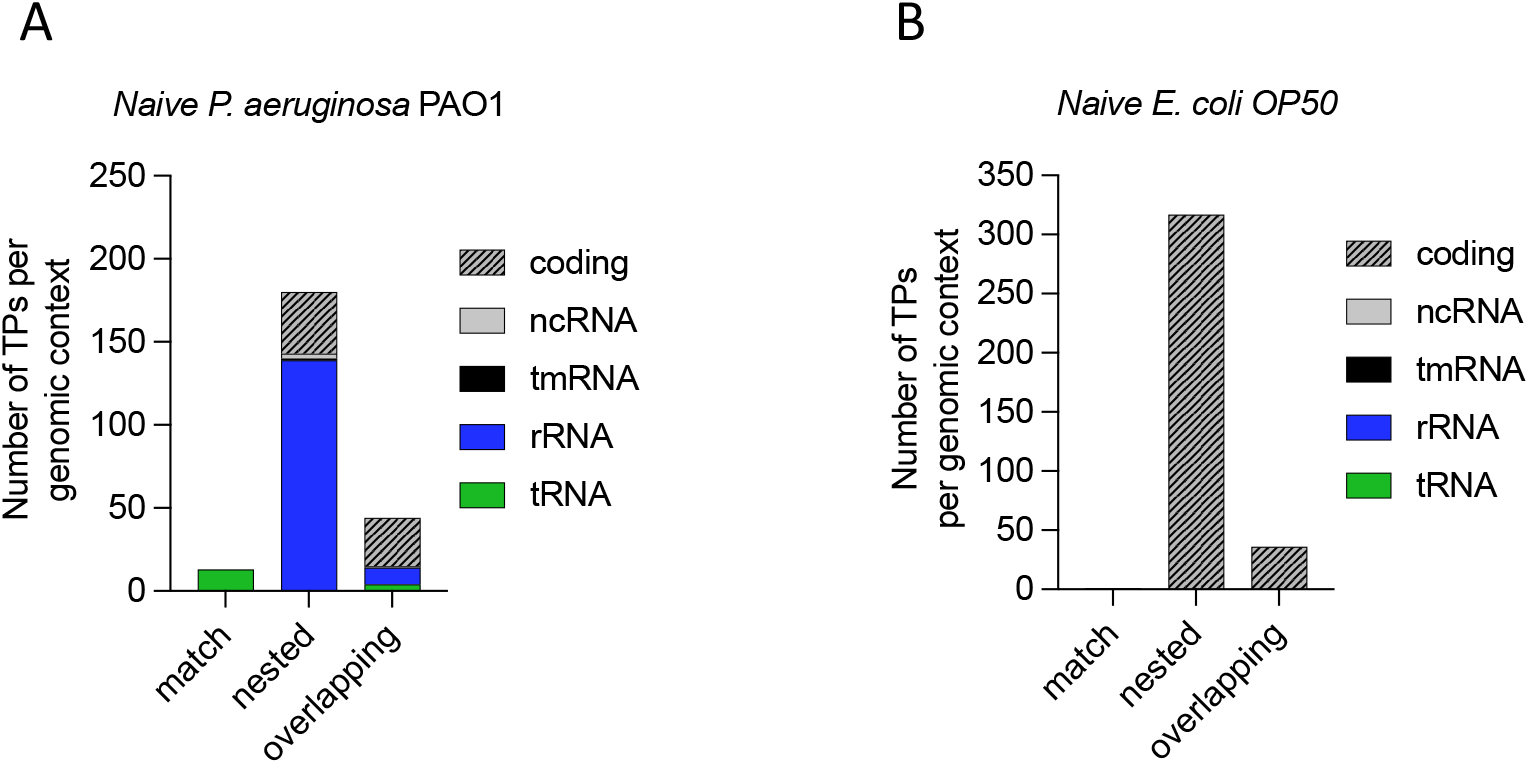
**A-B** Number of normalized TPs of naïve *P. aeruginosa* (A) and *E. coli* (B) per genomic context on each RNA biotype.

Table S1. Full list of TPs and their genomic contexts.

Table S2. RsmY complementary hits in the *C. elegans* genome.

File S1. Differentially expressed genes in *P. aeruginosa* PAO1 in the transgenerational paradigm in the intestine of F1, F2, F5 and F6 *C. elegans* compared to naïve.

File S2. Differentially expressed genes in *E. coli* OP50 in the transgenerational paradigm in the intestine of F3 and F4 *C. elegans* compared to naïve.

File S3. Differentially expressed genes in *C. elegans* in the transgenerational paradigm for six generations

(F1, F2, F5 and F6 feeding in *P. aeruginosa* PAO1; F3 and F4 feeding in *E. coli* OP50)

Source Data File 1. All data in the manuscript and statistical analysis.

## Acknowledgments

This work was done under extraordinary circumstances far beyond the COVID pandemic. We are deeply grateful to Marcia Manterola who provided a laboratory space in the later stages of the investigation and Ramon Latorre who gave unconditional support. Tatiana Leo allowed AC to be in two places at the same time. We thank Chiayu Chiu and Irini Topalidou for critical reading of the manuscript and Carolaing Gabaldon for technical contributions. Some strains were provided by the CGC, which is funded by NIH Office of Research Infrastructure Programs (P40OD010440). Dr. Stephan Heeb from the University of Nottingham provided the *P. aeruginosa* PAO1-L wild type and mutant strains. Dr. Meng Wang from the Baylor College of Medicine provided the *modified RNAi-E. coli OP50* strain.

## Author contribution

Conceptualization ML and AC

Methodology ML, BP and AC

Investigation ML, BP, BA and AC

Writing-Original Draft ML and AC

Writing-Review and Editing ML and AC

Funding Acquisition AC

## References

Aballay, A., & Hong, C. (2020). Microbial colonization induces histone acetylation critical for inherited gut-germline-neural signaling. bioRxiv. Retrieved from https://www.biorxiv.org/content/biorxiv/early/2020/09/11/2020.09.10.291823.full.pdf

Alegado, R. A., & King, N. (2014). Bacterial influences on animal origins. Cold Spring Harb Perspect Biol, 6(11), a016162. doi:10.1101/cshperspect.a016162

Anders, S., & Huber, W. (2010). Differential expression analysis for sequence count data. Genome Biol, 11(10), R106. doi:10.1186/gb-2010-11-10-r106

Anders, S., Pyl, P. T., & Huber, W. (2015). HTSeq--a Python framework to work with high-throughput sequencing data. Bioinformatics, 31(2), 166–169. doi:10.1093/bioinformatics/btu638

Betin, V., Penaranda, C., Bandyopadhyay, N., Yang, R., Abitua, A., Bhattacharyya, R. P.,… Hung, D. T. (2019). Hybridization-based capture of pathogen mRNA enables paired host-pathogen transcriptional analysis. Sci Rep, 9(1), 19244. doi:10.1038/s41598-019-55633-6

Brencic, A., McFarland, K. A., McManus, H. R., Castang, S., Mogno, I., Dove, S. L., & Lory, S. (2009). The GacS/GacA signal transduction system of Pseudomonas aeruginosa acts exclusively through its control over the transcription of the RsmY and RsmZ regulatory small RNAs. Mol Microbiol, 73(3), 434–445. doi:10.1111/j.1365-2958.2009.06782.x

Brenner, S. (1974). The genetics of Caenorhabditis elegans. Genetics, 77(1), 71–94. Retrieved from https://www.ncbi.nlm.nih.gov/pubmed/4366476

Camacho, C., Coulouris, G., Avagyan, V., Ma, N., Papadopoulos, J., Bealer, K., & Madden, T. L. (2009). BLAST+: architecture and applications. BMC Bioinformatics, 10, 421. doi:10.1186/1471-2105-10-421

Cassada, R. C., & Russell, R. L. (1975). The dauerlarva, a post-embryonic developmental variant of the nematode Caenorhabditis elegans. Dev Biol, 46(2), 326–342. doi:10.1016/0012-1606(75)90109-8

Celluzzi, A., & Masotti, A. (2016). How Our Other Genome Controls Our Epi-Genome. Trends Microbiol, 24(10), 777–787. doi:10.1016/j.tim.2016.05.005

Chávez, F. P., & Calixto, A. (2019). Use of C. elegans Diapause to Study Transgenerational Responses to Pathogen Infection. Methods Mol Biol, 1918, 191–197. doi:10.1007/978-1-4939-9000-9_16

Cobb, N. A. (1914). Nematodes and their relationships. US Government Printing Office. Retrieved from https://pdfs.semanticscholar.org/e68d/89a1fc3487652a11be78a1ecae8790d196a9.pdf

Devanapally, S., Ravikumar, S., & Jose, A. M. (2015). Double-stranded RNA made in C. elegans neurons can enter the germline and cause transgenerational gene silencing. Proc Natl Acad Sci U S A, 112(7), 2133–2138. doi:10.1073/pnas.1423333112

Diallo, I., & Provost, P. (2020). RNA-Sequencing Analyses of Small Bacterial RNAs and their Emergence as Virulence Factors in Host-Pathogen Interactions. Int J Mol Sci, 21(5). doi:10.3390/ijms21051627

Fire, A., Xu, S., Montgomery, M. K., Kostas, S. A., Driver, S. E., & Mello, C. C. (1998). Potent and specific genetic interference by double-stranded RNA in Caenorhabditis elegans. Nature, 391(6669), 806–811. doi:10.1038/35888

Gabaldón, C., Legüe, M., Palominos, M. F., Verdugo, L., Gutzwiller, F., & Calixto, A. (2020). Intergenerational Pathogen-Induced Diapause in Caenorhabditis elegans Is Modulated by mir-243. mBio, 11(5). doi:10.1128/mBio.01950-20

Gu, H., Zhao, C., Zhang, T., Liang, H., Wang, X. M., Pan, Y.,… Zen, K. (2017). Salmonella produce microRNA-like RNA fragment Sal-1 in the infected cells to facilitate intracellular survival. Sci Rep, 7(1), 2392. doi:10.1038/s41598-017-02669-1

Heurlier, K., Williams, F., Heeb, S., Dormond, C., Pessi, G., Singer, D.,… Haas, D. (2004). Positive control of swarming, rhamnolipid synthesis, and lipase production by the posttranscriptional RsmA/RsmZ system in Pseudomonas aeruginosa PAO1. J Bacteriol, 186(10), 2936–2945. doi:10.1128/jb.186.10.2936-2945.2004

Houri-Ze’evi, L., & Rechavi, O. (2016). Plastic germline reprogramming of heritable small RNAs enables maintenance or erasure of epigenetic memories. RNA Biol, 13(12), 1212–1217. doi:10.1080/15476286.2016.1229732

Houri-Zeevi, L., & Rechavi, O. (2017). A Matter of Time: Small RNAs Regulate the Duration of Epigenetic Inheritance. Trends Genet, 33(1), 46–57. doi:10.1016/j.tig.2016.11.001

Jablonka, E., & Raz, G. (2009). Transgenerational epigenetic inheritance: prevalence, mechanisms, and implications for the study of heredity and evolution. Q Rev Biol, 84(2), 131–176. Retrieved from https://www.ncbi.nlm.nih.gov/pubmed/19606595

Jose, A. M. (2015). Movement of regulatory RNA between animal cells. Genesis, 53(7), 395–416. doi:10.1002/dvg.22871

Kaletsky, R., Moore, R. S., Vrla, G. D., Parsons, L. R., Gitai, Z., & Murphy, C. T. (2020). C. elegans interprets bacterial non-coding RNAs to learn pathogenic avoidance. Nature. doi:10.1038/s41586-020-2699-5

Kalinava, N., Ni, J. Z., Peterman, K., Chen, E., & Gu, S. G. (2017). Decoupling the downstream effects of germline nuclear RNAi reveals that H3K9me3 is dispensable for heritable RNAi and the maintenance of endogenous siRNA-mediated transcriptional silencing in. Epigenetics Chromatin, 10, 6. doi:10.1186/s13072-017-0114-8

Kawli, T., & Tan, M. W. (2008). Neuroendocrine signals modulate the innate immunity of Caenorhabditis elegans through insulin signaling. Nat Immunol, 9(12), 1415–1424. doi:10.1038/ni.1672

Kay, E., Humair, B., Dénervaud, V., Riedel, K., Spahr, S., Eberl, L.,… Haas, D. (2006). Two GacA-dependent small RNAs modulate the quorum-sensing response in Pseudomonas aeruginosa. J Bacteriol, 188(16), 6026–6033. doi:10.1128/JB.00409-06

Koeppen, K., Hampton, T. H., Jarek, M., Scharfe, M., Gerber, S. A., Mielcarz, D. W.,… Stanton, B. A. (2016). A Novel Mechanism of Host-Pathogen Interaction through sRNA in Bacterial Outer Membrane Vesicles. PLoS Pathog, 12(6), e1005672. doi:10.1371/journal.ppat.1005672

Langmead, B., & Salzberg, S. L. (2012). Fast gapped-read alignment with Bowtie 2. Nat Methods, 9(4), 357–359. doi:10.1038/nmeth.1923

Legüe, M., & Calixto, A. (2019). RNA language in Caenorhabditis elegans and bacteria interspecies communication and memory. Current Opinion in Systems Biology, 13. Retrieved from https://doi.org/10.1016/j.coisb.2018.08.005

Leitão, A. L., Costa, M. C., Gabriel, A. F., & Enguita, F. J. (2020). Interspecies Communication in Holobionts by Non-Coding RNA Exchange. Int J Mol Sci, 21(7). doi:10.3390/ijms21072333

Lev, I., Seroussi, U., Gingold, H., Bril, R., Anava, S., & Rechavi, O. (2017). MET-2-Dependent H3K9 Methylation Suppresses Transgenerational Small RNA Inheritance. Curr Biol, 27(8), 1138–1147. doi:10.1016/j.cub.2017.03.008

Liao, Y., Smyth, G. K., & Shi, W. (2014). featureCounts: an efficient general purpose program for assigning sequence reads to genomic features. Bioinformatics, 30(7), 923–930. doi:10.1093/bioinformatics/btt656

Liu, S., da Cunha, A. P., Rezende, R. M., Cialic, R., Wei, Z., Bry, L.,… Weiner, H. L. (2016). The Host Shapes the Gut Microbiota via Fecal MicroRNA. Cell Host Microbe, 19(1), 32–43. doi:10.1016/j.chom.2015.12.005

Liu, W., Rochat, T., Toffano-Nioche, C., Le Lam, T. N., Bouloc, P., & Morvan, C. (2018). Assessment of Bona Fide sRNAs in Staphylococcus aureus. Front Microbiol, 9, 228. doi:10.3389/fmicb.2018.00228

Love, M. I., Huber, W., & Anders, S. (2014). Moderated estimation of fold change and dispersion for RNA-seq data with DESeq2. Genome Biol, 15(12), 550. doi:10.1186/s13059-014-0550-8

Low, L. Y., Harrison, P. F., Gould, J., Powell, D. R., Choo, J. M., Forster, S. C.,… Rood, J. I. (2018). Concurrent Host-Pathogen Transcriptional Responses in a Clostridium perfringens Murine Myonecrosis Infection. mBio, 9(2). doi:10.1128/mBio.00473-18

Lybecker, M., Bilusic, I., & Raghavan, R. (2014). Pervasive transcription: detecting functional RNAs in bacteria. Transcription, 5(4), e944039. doi:10.4161/21541272.2014.944039

Mao, H., Zhu, C., Zong, D., Weng, C., Yang, X., Huang, H.,… Guang, S. (2015). The Nrde Pathway Mediates Small-RNA-Directed Histone H3 Lysine 27 Trimethylation in Caenorhabditis elegans. Curr Biol, 25(18), 2398–2403. doi:10.1016/j.cub.2015.07.051

Marré, J., Traver, E. C., & Jose, A. M. (2016). Extracellular RNA is transported from one generation to the next in Caenorhabditis elegans. Proc Natl Acad Sci U S A, 113(44), 12496–12501. doi:10.1073/pnas.1608959113

Martin, M. (2011). Cutadapt removes adapter sequences from high-throughput sequencing reads. EMBnet. journal, 17(1), 10–12. Retrieved from https://journal.embnet.org/index.php/embnetjournal/article/viewFile/200/458

Moore, R. S., Kaletsky, R., & Murphy, C. T. (2019). Piwi/PRG-1 Argonaute and TGF-β Mediate Transgenerational Learned Pathogenic Avoidance. Cell, 177(7), 1827–1841.e12. doi:10.1016/j.cell.2019.05.024

Nono, M., Kishimoto, S., Sato-Carlton, A., Carlton, P. M., Nishida, E., & Uno, M. (2020). Intestine-to-Germline Transmission of Epigenetic Information Intergenerationally Ensures Systemic Stress Resistance in C. elegans. Cell Rep, 30(10), 3207–3217.e4. doi:10.1016/j.celrep.2020.02.050

Palominos, M. F., Verdugo, L., Gabaldon, C., Pollak, B., Ortíz-Severín, J., Varas, M. A.,… Calixto, A. (2017). Transgenerational Diapause as an Avoidance Strategy against Bacterial Pathogens in Caenorhabditis elegans. mBio, 8(5). doi:10.1128/mBio.01234-17

Pereira, A. G., Gracida, X., Kagias, K., & Zhang, Y. (2020). <i>C. elegans</i> aversive olfactory learning generates diverse intergenerational effects. J Neurogenet, 1–11. doi:10.1080/01677063.2020.1819265

Pessi, G., Williams, F., Hindle, Z., Heurlier, K., Holden, M. T., Cámara, M.,… Williams, P. (2001). The global posttranscriptional regulator RsmA modulates production of virulence determinants and N-acylhomoserine lactones in Pseudomonas aeruginosa. J Bacteriol, 183(22), 6676–6683. doi:10.1128/JB.183.22.6676-6683.2001

Poinar, G. O. (2015). The Geological Record of Parasitic Nematode Evolution. Adv Parasitol, 90, 53–92. doi:10.1016/bs.apar.2015.03.002

Posner, R., Toker, I. A., Antonova, O., Star, E., Anava, S., Azmon, E.,… Rechavi, O. (2019). Neuronal Small RNAs Control Behavior Transgenerationally. Cell, 177(7), 1814–1826.e15. doi:10.1016/j.cell.2019.04.029

Provorov, N. A., Vorob’ev, N. I., & Andronov, E. E. (2008). [Macro- and microevolution of bacteria in symbiotic systems]. Genetika, 44(1), 12–28. Retrieved from https://www.ncbi.nlm.nih.gov/pubmed/18409383

Quinlan, A. R., & Hall, I. M. (2010). BEDTools: a flexible suite of utilities for comparing genomic features. Bioinformatics, 26(6), 841–842. doi:10.1093/bioinformatics/btq033

Rechavi, O., Houri-Ze’evi, L., Anava, S., Goh, W. S. S., Kerk, S. Y., Hannon, G. J., & Hobert, O. (2014). Starvation-induced transgenerational inheritance of small RNAs in C. elegans. Cell, 158(2), 277–287. doi:10.1016/j.cell.2014.06.020

Rechavi, O., & Lev, I. (2017). Principles of Transgenerational Small RNA Inheritance in Caenorhabditis elegans. Curr Biol, 27(14), R720–R730. doi:10.1016/j.cub.2017.05.043

Remy, J. J. (2010). Stable inheritance of an acquired behavior in Caenorhabditis elegans. Curr Biol, 20, R877–8. doi:10.1016/j.cub.2010.08.013

Robinson, M. D., McCarthy, D. J., & Smyth, G. K. (2010). edgeR: a Bioconductor package for differential expression analysis of digital gene expression data. Bioinformatics, 26(1), 139–140. doi:10.1093/bioinformatics/btp616

Robinson, M. D., & Oshlack, A. (2010). A scaling normalization method for differential expression analysis of RNA-seq data. Genome Biol, 11(3), R25. doi:10.1186/gb-2010-11-3-r25

Rutherford, S. T., & Bassler, B. L. (2012). Bacterial quorum sensing: its role in virulence and possibilities for its control. Cold Spring Harb Perspect Med, 2(11). doi:10.1101/cshperspect.a012427

Samuel, B. S., Rowedder, H., Braendle, C., Félix, M. A., & Ruvkun, G. (2016). Caenorhabditis elegans responses to bacteria from its natural habitats. Proc Natl Acad Sci U S A, 113(27) E3941–9. doi:10.1073/pnas.1607183113

Skinner, M. K. (2011). Environmental epigenetic transgenerational inheritance and somatic epigenetic mitotic stability. Epigenetics, 6(7), 838–842. Retrieved from https://www.ncbi.nlm.nih.gov/pubmed/21637037

Stiernagle, T. (2006). Maintenance of C. elegans. WormBook, 1–11. doi:10.1895/wormbook.1.101.1

Subramanian, D., Bhasuran, B., & Natarajan, J. (2019). Genomic analysis of RNA-Seq and sRNA-Seq data identifies potential regulatory sRNAs and their functional roles in Staphylococcus aureus. Genomics, 111(6), 1431–1446. doi:10.1016/j.ygeno.2018.09.016

Valverde, C., Heeb, S., Keel, C., & Haas, D. (2003). RsmY, a small regulatory RNA, is required in concert with RsmZ for GacA-dependent expression of biocontrol traits in Pseudomonas fluorescens CHA0. Mol Microbiol, 50(4), 1361–1379. doi:10.1046/j.1365-2958.2003.03774.x

van den Hoogen, J., Geisen, S., Routh, D., Ferris, H., Traunspurger, W., Wardle, D. A.,… Crowther, T. W. (2019). Soil nematode abundance and functional group composition at a global scale. Nature, 572(7768), 194–198. doi:10.1038/s41586-019-1418-6

Westermann, A. J., Förstner, K. U., Amman, F., Barquist, L., Chao, Y., Schulte, L. N.,… Vogel, J. (2016). Dual RNA-seq unveils noncoding RNA functions in host-pathogen interactions. Nature, 529(7587), 496–501. doi:10.1038/nature16547

Westermann, A. J., Gorski, S. A., & Vogel, J. (2012). Dual RNA-seq of pathogen and host. Nat Rev Microbiol, 10(9), 618–630. doi:10.1038/nrmicro2852

Winsor, G. L., Griffiths, E. J., Lo, R., Dhillon, B. K., Shay, J. A., & Brinkman, F. S. (2016). Enhanced annotations and features for comparing thousands of Pseudomonas genomes in the Pseudomonas genome database. Nucleic Acids Res, 44(D1), D646–53. doi:10.1093/nar/gkv1227

Woodhouse, R. M., & Ashe, A. (2019). Transgenerational Epigenetic Inheritance Is Revealed as a Multi-step Process by Studies of the SET-Domain Proteins SET-25 and SET-32. Epigenet Insights, 12, 2516865719844214. doi:10.1177/2516865719844214

Xiao, R., Chun, L., Ronan, E. A., Friedman, D. I., Liu, J., & Xu, X. Z. (2015). RNAi Interrogation of Dietary Modulation of Development, Metabolism, Behavior, and Aging in C. elegans. Cell Rep, 11(7), 1123–1133. doi:10.1016/j.celrep.2015.04.024

Zhang, Y., Lu, H., & Bargmann, C. I. (2005). Pathogenic bacteria induce aversive olfactory learning in Caenorhabditis elegans. Nature, 438(7065), 179–184. doi:10.1038/nature04216

Zhao, C., Zhou, Z., Zhang, T., Liu, F., Zhang, C. Y., Zen, K., & Gu, H. (2017). Salmonella small RNA fragment Sal-1 facilitates bacterial survival in infected cells via suppressing iNOS induction in a microRNA manner. Sci Rep, 7(1), 16979. doi:10.1038/s41598-017-17205-4

